# Reflection spectroscopy of bistable visual pigments in living butterflies

**DOI:** 10.64898/2026.05.15.725499

**Authors:** Primož Pirih

## Abstract

Invertebrate vision relies on bistable visual pigments flipping upon photon absorption between rhodopsin and metarhodopsin states. In living butterflies, the UV-VIS absorption spectra of rhodopsin and metarhodopsin, respectively with 11-cis and all-trans isomers of 3-hydroxy-retinal (A3) chromophore, can be conveniently recorded from the eyeshine, the light reflected from the compound eye after passing twice through the light-guiding rhabdoms. **\*** Here, a microscope coupled with a broadband LED source and a microspectrometer was used to record photorelaxations reported in eyeshine reflection spectra. Fitting temporal exponential relaxations to log-reflectance arrays yielded transient and baseline spectra that are analogous to absorbance difference and sum, respectively. Both types of spectra were subjected to singular value decomposition and to fitting of templated visual pigment absorption spectra. **\*** The compound eye of the high brown fritillary *Fabriciana adippe* was exposed to a series of second-long broadband light pulses, causing photorelaxations with time constants between 40 and 120 ms that led to 80% metarhodopsin in equilibrium. The transient and baseline spectra were fitted with pigment templates, estimating the alpha peak wavelength 547-552 nm for rhodopsin and 496-501 nm for metarhodopsin. The metarhodopsin to rhodopsin alpha peak absorbance ratio 1.25-1.35 is consistent with the isosbestic wavelength at 530 nm. The second isosbestic wavelength indicates that rhodopsin beta (UV) peak absorbs more strongly than metarhodopsin below 405 nm. **\*** Baseline spectra, which were not explicitly analysed in previous studies, enable concatenation of exposures, monitor long-term changes of pigment, and enhance the estimation of beta peak parameters. **\*** The method can be directly used in many butterflies and could be adapted to other insects, particularly fruitflies, facilitating studies of the relation between the visual pigment spectra and the opsin sequences. Spectroscopic results can be complemented with physiologically measured photoreceptor spectral sensitivity datasets and analysed with the same global fitting procedure.

## 1 Introduction

The compound eye of butterflies consists of several thousand anatomically similar ommatidia. Each ommatidium contains nine photoreceptor cells (Fig. 1). Their photoreceptive organelles, the rhabdomeres, are closely packed microvilli that contain visual pigment molecules. The rhabdomeres form together a fused rhabdom, which acts as a light guide. In nymphalid butterflies, the receptors R1-2 have relatively short rhabdomeres located in the distal tier of the ommatidium and contain UV-or blue-peaking rhodopsins. Photoreceptors R3-R8 that contribute to the rhabdom along its whole length, and the basal R9, contain the main (long wavelength, LW) visual pigment, whose rhodopsin form absorbs maximally in the wavelength range 500 .. 600 nm [^1,6,14,31,45,54^].

**Figure 1.**
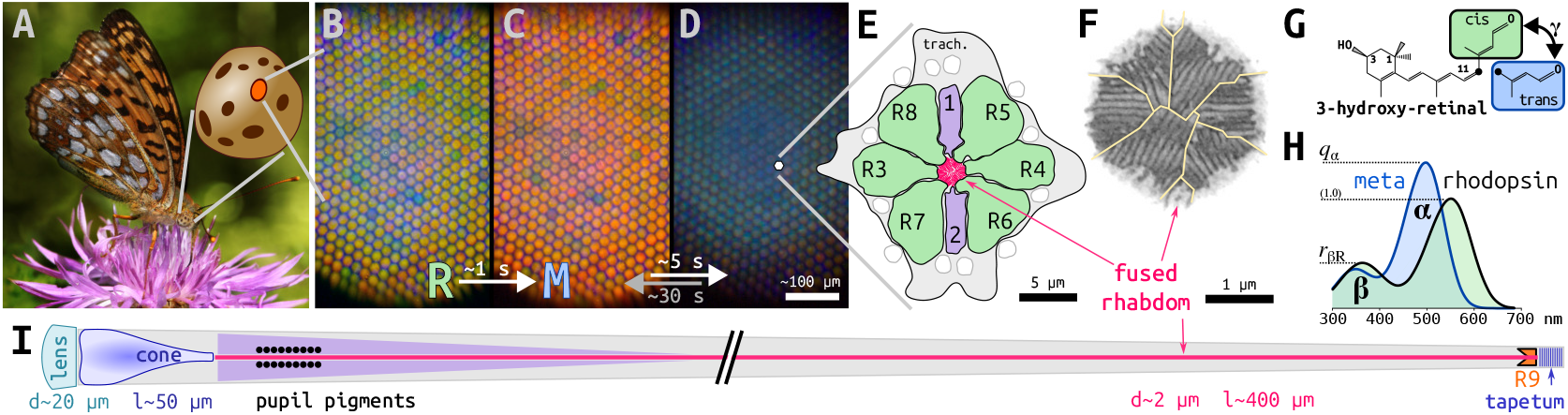
The eye of a brushfoot butterfly. **(A)** The high brown fritillary *Fabriciana adippe*, [photo by Richard Bartz ^4^] **(B)** The dark-adapted female eyeshine (*simulated*), **(C)** after photorelaxation, **(D)** after pupil closure. **(E)** Scheme of a cross-section of an ommatidium. The microvilli of photoreceptors R1-R2 (*lavender*) and R3-R8 (*jade*) form a fused rhabdom (*magenta*). The photoreceptors are surrounded by glial cells (*gray*). **(F)** Electron micrograph of the rhabdom in the proximal eye part (courtesy of prof. Kentaro Arikawa). At this depth, the rhabdomeres of the photoreceptors R3-R8 contribute to the fused rhabdom. **(G)** Butterfly visual pigment contains 3-hydroxy-retinal (A3 chromophore), in 11-cis configuration in *rhodopsin* (*jade*), in all-trans configuration in *metarhodopsin* (*sky*). **(H)** LW metarhodopsin is *hypsochromic* (blue-shifted) and has higher alpha peak absorption than rhodposin. The ratio of peak absorptions is *q*_*α*_ *>* 1. **(I)** Longitudinal scheme of an ommatidium (*not to scale*). Light is being launched through the distal optics (*left: lens, cone*) into the rhabdom waveguide (*magenta*) and is reflected back by the *tapetum* multilayer at the bottom. Photoreceptors R1-2 (*lavender*) contribute to the rhabdom distally, R9 (*orange*) basally.

A multilayered tapetum is situated below each of the light-guiding rhabdoms [^51,60^]. The tapetum plays a crucial role in the optical physiology of butterfly eyes. Incident light is focussed by the facet lens and the crystalline cone, guided by the rhabdom, and back-reflected by the tapetum (Fig. 1I). The light not absorbed by visual and screening pigments on the downwelling and upwelling path radiates out as the eyeshine [^74,75,84^]. A superimposed image of ommatidial reflections forms a bright spot – the luminous pseudopupil – in the centre of the eye spheroid. This unique feature can be exploited for spectroscopical studies of visual pigment photochemistry [^11,87^] and for measurements of pupillary optophysiology [^57,58^].

The microvillar membranes of the rhabdomeres contain visual pigments, trans-membrane proteins (opsins) bound with a retinal chromophore. In contrast to vertebrate visual pigments, where the metarhodopsin state rapidly degrades (bleaches) to opsin and chromophore, invertebrate visual pigments are *bistable*, that is, their metarhodopsin is relatively long-lived [^80,86^]. Upon absorption of a photon, the invertebrate rhodopsin (R, with 11-cis chromophore) passes through short-lived intermediates [^33^], reaching metarhodopsin state (M, with all-trans chromophore) that activates the Gq-protein cascade. This leads to an influx of sodium and calcium through TRP channels, resulting in cell membrane depolarisation [^35,41,49^]. Heightened intracellular calcium causes the granules of pupillary pigment to closely encircle the rhabdom, absorbing a large fraction of light [^44,45^]. In butterflies, the pupil usually starts closing within a second after the onset of illumination [^57,81^]. The phototransducing form of metarhodopsin is either reconverted back to rhodopsin by absorbing another photon, or deactivated by binding to arrestin and phosphorylation, followed by removal from the rhabdomere and degradation in the soma (bleaching) [^5,65,71,79^]. In butterflies, removal and resynthesis operate on quarter-hourly and hourly scale, respectively, dependent on the temperature [^9,10,57,87^].

Visual pigments have two physiologically relevant absorption peaks, the alpha and the beta (UV) peak (1F). Their absorption spectra can be approximated by normalised *templates*, parametrised by the alpha peak position *λ*_*α*_ [^28,32,77,83^]. The alpha peak magnitude of metarhodopsin is higher than that of rhodopsin, their ratio is reported in the range (1.2 *< q*_*α*_ *<* 2.0) [^80^]. Flies and butterflies have A3 visual pigments with 3-hydroxyretinal as the chromophore [^30,66,89^]. The A3 rhodopsin beta to alpha peak absorption ratio is assumed to be *r*_*β*R_ = 0.25, as in vertebrate A1 and A2 rhodopsins [^32,56^], although electrophysiological data often hints that this value can be different [^42,58,68^]. The LW rhodopsin of butterflies has the alpha peak in the range [500 nm *< λ*_*α*R_ *<* 600 nm]. The alpha peak of its metarhodopsin is blue-shifted (hypsochromic), peaking in the range [480 nm *< λ*_*α*M_ *<* 510 nm] (Fig. 1H). The exact metarhodopsin absorption spectra are less well known, but it is assumed that they follow the templates for A1 rhodopsin; a correction for the fly A3 metarhodopsin template was given by [^77^].

Butterfly eyeshine serves as a convenient platform that enables *in vivo* spectroscopic studies of visual pigments. Spectroscopy has been conducted using low intensity monochromatic scanning light that caused negligible photoconversion and pupil activation [^8–11,13,21,26,27^]. An alternative is to use an epi-illumination microscope system with a dispersive UV-VIS spectrometer and a broadband light source that causes the relaxation of rhodopsin-metarhodopsin photosystem to be completed within a fraction of a second, before the activation of the pupil. Exposure to broadband light causes a relaxation process that can be approximated by an exponential function. The measured change in the logarithmically-transformed reflection spectra is proportional to the difference between rhodopsin and metarhodopsin absorption spectra [^87^]. The accuracy of rhodopsin absorption spectra estimated with broadband spectroscopy has been validated through comparison with sensitivity spectra obtained with intracellular electrophysiology and pupil optical retinography (ORG) [^58^]. Butterfly visual pigments have also been spectroscopically studied in cell expression systems [^21,27,47,63,93^], and expressed in the fruitfly and studied with pupillary optophysiology [^27^] and extracellular electrophysiology [^68^].

Outside butterflies, photochemistry of invertebrate bistable visual pigments has been studied using UV-VIS absorption spectroscopy in intact fruitfly [^72^] and blowfly eyes [^46,61^], using antidromic (backward) illumination. The more involved preparations include slices [^19,20,29,53,64^], purified crystals [^52,88^] and extracts [^24,64,92^]. Fundamental studies are reviewed in [^80^].

Here, a method for acquisition and comprehensive analysis of broadband UV-VIS eyeshine reflectance spectra is described. Arrays of reflectance spectra were acquired in one second exposures, interleaved with longer (20–140 min) dark intervals. Each log-reflectance array was fitted with an exponential relaxation model, yielding two spectra: the transient spectrum described the photochemical relaxation completed within a fraction of a second, the baseline spectrum monitored changes that were due to the synthesis and removal (bleaching) of the visual pigment on hourly scale. The two spectra are analogous to double-pass absorbance difference and sum, respectively. The spectra were subjected to singular value decomposition (SVD) and to fitting with pigment absorption template spectra. Inclusion of baseline spectra improved the analysis, particularly with respect to the estimation of beta peaks. Modifications of the fitting procedure to account for non-linearities and rhabdom waveguide effects, and adaptations of the spectroscopic method to other insect groups are discussed.

## 2 Experimental method

The female high brown fritillaries *Fabriciana adippe* (Nymphalidae: Heliconinae: Argynnini) were collected around Ljubljana, Slovenia. The butterflies were immobilized into a cut pipette tip or a thermal insulation tube using a mixture of beeswax and colophony, and placed into a rotational stage under a modified Leitz Orthoplan microscope equipped with an epi-illumination module with a 50% beam splitter and Olympus MPlan 10 × 0.25(0*/*∞) objective. The spectrometer head had a circular diaphragm at the first focal plane, a side viewing port, and two lenses that relayed the diaphragm plane to the rosette-to-linear fibre bundle (BFL200HS02, Thorlabs) connected to the microspectrometer (USB2000, Ocean Insight) controlled by a Raspberry PI 4 computer. The light source was a white phosphorescent LED based on a violet emitter (Soraa MR16-50-B03, 405 nm, modified for DC current). The useful spectroscopic range of about 390–800 nm was estimated from a piece of pressed magnesium oxide powder. The shutter (SHBH1, Thorlabs) was controlled by the spectrometer; electronic switching could not be used due to thermal effects on the LED emission spectrum. Eyeshine images were taken by replacing the fibre head with a C-mount camera (Raspberry HQ camera with the NIR–blocking filter removed, IMX477, 4056*3040 pixels, 1.55 µm pitch). The deep luminous pseudopupil was focussed and centred on the diaphragm by translating the stage while being observed through a telescopic eyepiece and the spectrometer head’s side viewing port. Spectrometer signal was finally optimized in a Python-based GUI [^15,59^] by focussing the final lens and laterally translating the fibre holder.

The spectra were acquired in GNU Octave through a utility written for C-Seabreeze library, and analysed in GNU Octave with packages Signal and Optim [versions 9.3.0, 3.0.11; 11.1.0, 1.4.6, 1.6.2, respectively; ^23,55^]. The integration time was set to 11 ms in order to minimise the USB bus jitter. The spectrometer’s 12-bit analogue-digital converter (ADC) had offset value (code) about 200. The offset was corrected by subtracting the average of 100 spectra taken immediately before each exposure. The maximal ADC value converted to decadic logarithm was thus log_10_ 3900 ≈ 3.6. Prior to exposures, two optical dark reference spectra were measured with the same integration time and additional averaging. The pupil reference was recorded with the eye in the experimental position after the shutter was open for one minute to cause maximal pupil closure. Because the closed pupil leaked some light in the blue spectral range (Fig. 1C), the animal was moved sideways so that the eyeshine was blocked by the aperture. The side reference was scaled down to match the pupil reference in the long wavelength part of the spectrum and used as the final optical dark reference.

## 3 Results

### 3.1 Relaxation model yields transient and baseline spectra

The spectroscopic data was acquired during one second exposures to broadband light. The 21 collected exposures were interleaved with dark intervals lasting between 20 and 140 minutes. In total, about 10 seconds of spectroscopic data was analysed. The array of spectra acquired in each exposure was dark reference-corrected, conditioned, rectified (Sec. 7.3) and log_10_-transformed. Each resulting array **L**_0_ was separately fitted with a first-order exponential relaxation model (Sec. 7.4). The model yielded a single non-linear parameter, the relaxation time constant *τ*, and two spectra – the transient spectrum **a** and the (raw) baseline spectrum **b**_0_ (Fig. 2B, *blue and magenta traces*). The transient spectra **a** obtained from 21 exposures (Fig. 2C) had similar shapes. Positive values of the transient amplitude spectrum **a** signify that the reflectance diminished during the exposure. The largest transients were acquired in the exposures following the longest dark intervals (*mauve traces*). The transient spectra passed the zero line twice, at the isosbestic wavelengths in the violet and green spectral range, *λ*_iso_ ≈ {406, 531 nm}.

**Figure 2.**
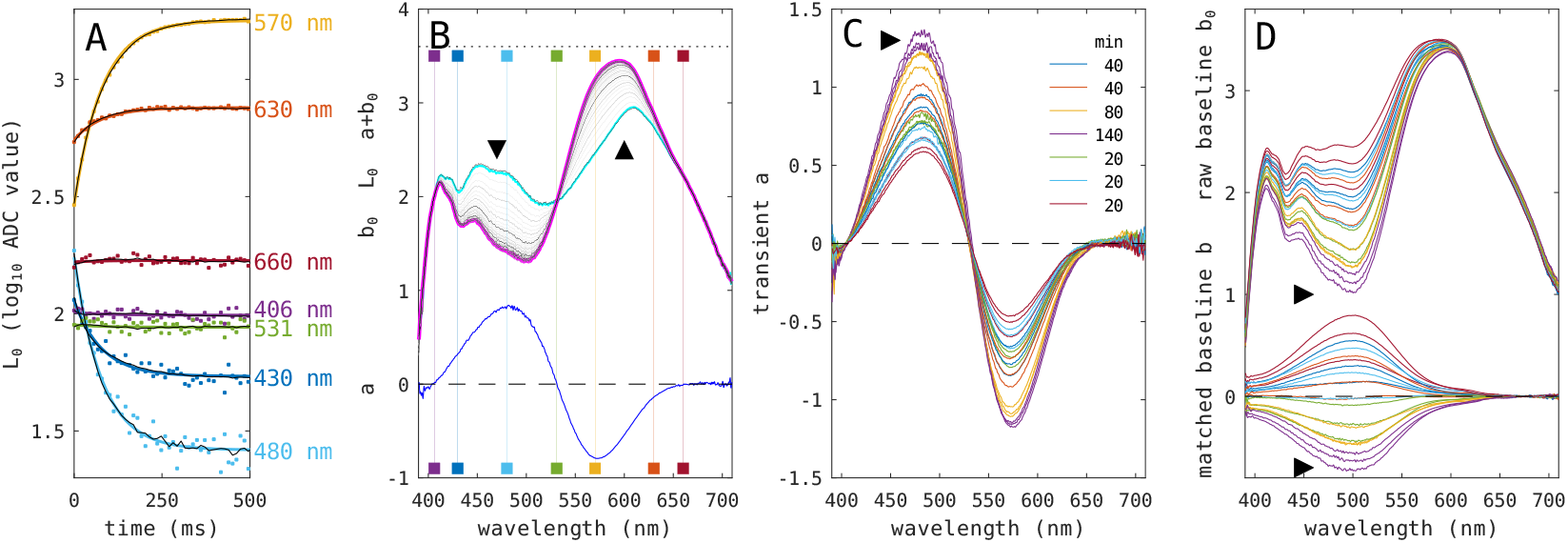
Transient and baseline spectra. **(A) Single exposure on the temporal axis**. Unconditioned (*coloured dots*) and conditioned traces (*black*). Exponential relaxation fit model at seven wavelengths (*coloured traces*). **(B) Single exposure on the spectral axis**. The log-transformed spectra **L**_0_ (*thin gray traces*). Transient amplitude spectrum (**a**) (blue trace), baseline spectrum **b**_0_ (*magenta trace*), starting spectrum (**a** + **b**_0_) (*cyan trace*). Direction of change (*arrows*), wavelengths shown in panel A (*squares*), maximal ADC value (*dashed line*). **(C) Transient spectra**. Preceding dark adaptation intervals (*legend*). **(D) Baseline spectra**. Raw baseline spectra **b**_0_ (*top part*), matched baseline spectra **b** (*bottom part*). *Right arrow* points to the traces with the highest noise.

Raw baseline spectra **b**_0_ (Fig. 2D, top) contain information about the quantity of visual pigments after the ommatidia have reached the photochemical equilibrium, but their shapes are modified by the experimental settings (light source, microscope transmittance, spectrometer sensitivity) and biological unknowns (cornea transmittance, tapetum reflectance). It is not trivial to measure the latter two, because a complete bleach of visual pigment is difficult to attain [^9,57^]. In order to exclude these effects, the baseline spectra were aligned in the red wavelength range and then had their ensemble average subtracted (Sec. 7.5). The resulting *matched* baseline spectra **b** exhibited a uniform shape (Fig. 2D, bottom). The spectra had more noise at both ends, and around 500 nm, particularly in the exposures obtained after long dark intervals (Fig. 2CD, *right arrow*).

### 3.2 Baseline correction enables concatenation of exposures

The absorption spectra of the main visual pigments are presumably not changing during the experimental time. Baseline correction enables seamless concatenation of the spectro-temporal arrays, thus increasing the spectroscopic information available to visual pigment template fitting procedure. The baseline-corrected arrays were obtained by subtracting their respective (raw) baseline spectra, **L** = **L**_0_ − **b**_0_ (Fig. 2C, top), and then concatenated to form a large array **L**_*C*_ *= [****L***_*[1:21]*_] (*Fig*. 3A, *lower part*). The array was subjected to singular value decomposition (SVD), yielding spectral component vectors 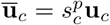 and temporal component vectors 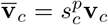 scaled with the square root of the component signal (*p* = 1*/*2) for display (Sec. 7.2). Temporal components 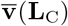 are shown in Fig. 3C. Spectral components **ū**(**L**_C_) are shown in Fig. 3D.

**Figure 3.**
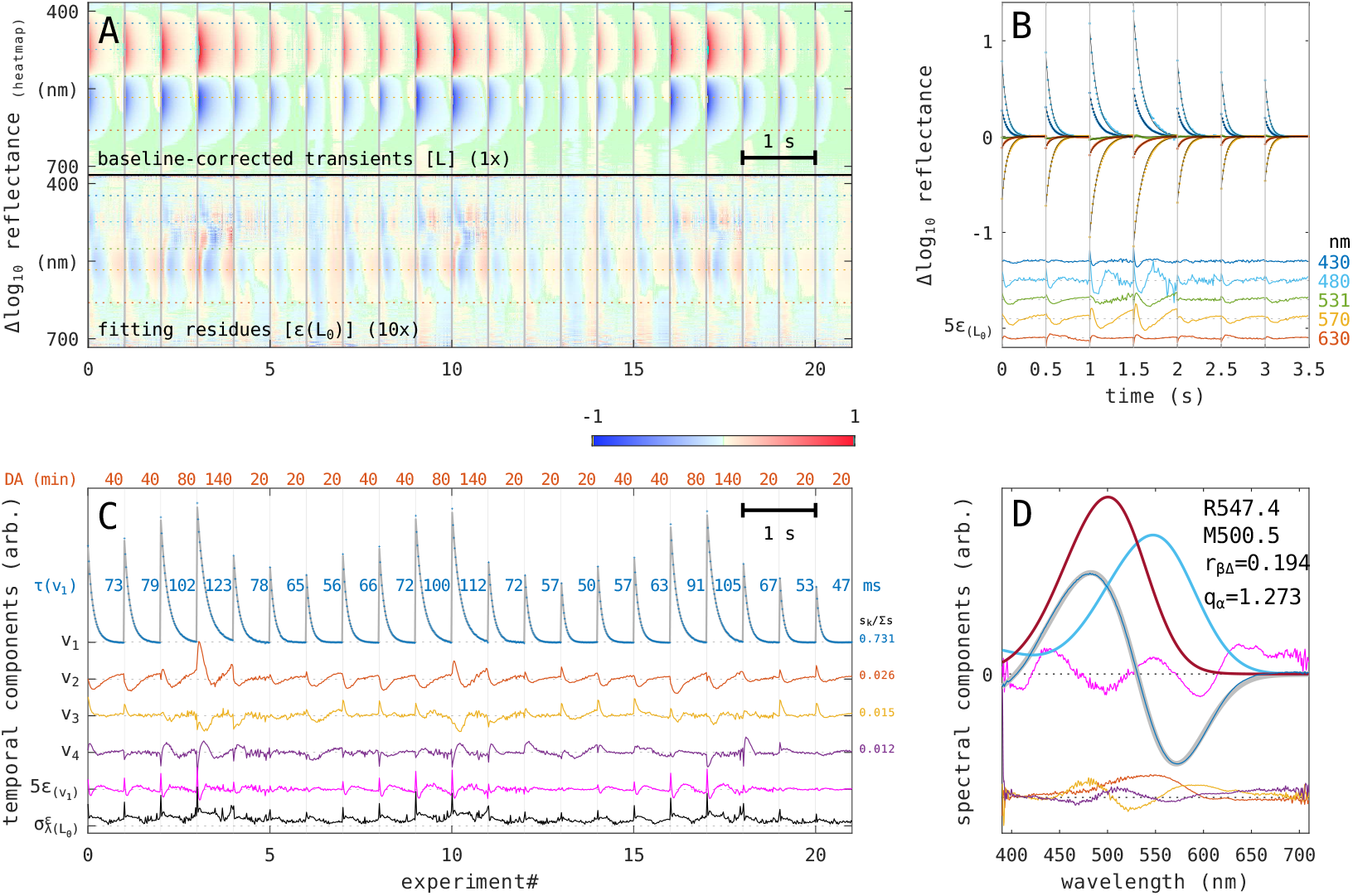
Concatenation of baseline-corrected transients. (**A**) **Baseline correction**. Concatenated heat map of the transients (**L**, *lower part*); the residues of fitting the exponential relaxation to individual arrays, *ϵ*(**L**_0_) (*upper part*, scaled 20*×*) **(B) Exemplary time traces**. Concatenated traces from seven exposures **L** at five wavelengths (*numbers right*), fitted exponential relaxations (*thin black traces*), fitting residues *ϵ*(**L**_0_) (*bottom, coloured traces*, scaled 5*×*). **(C) Temporal components from SVD of the concatenated array**. The components **ū**_1:4_(**L**_C_) shown with *blue, red, yellow, mauve traces*. First-order relaxation models fitted to **ū**_1_ (*gray trace*) and their residues *ϵ*(**ū**_1_) (*pink trace*, scaled 5*×*). Preceding dark adaptation intervals (*red*, minutes) and the estimated relaxation time constants (*blue*, milliseconds). The standard deviation of the residues *σ*_*λ*_(*ϵ*(**L**_0_)) (*black trace*, arbitrarily scaled, data from panel A, bottom). Relative component signals (*numbers right*). **(D) SVD spectral components and a template fit**. The first component **ū**_1_ (*blue trace*) fitted with a template model with constraint *r*_*β*M_ ≡ 0 (*thin black trace*). Components **ū**_2:4_ (offset; *red, yellow, mauve traces*). Rhodopsin template (*sky trace*), metarhodopsin template (*brown trace*), fit residuals *ϵ*(**ū**_1_) (*pink trace*, scaled 10*×*). Parameter estimates (*black text*).

The first temporal component 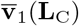 contained a sequence of photochemical relaxations with differing starting amplitudes and relaxation rates (*blue trace*). The first component contained about three quarters of the signal energy. The parts of 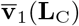 were piecewise fitted with the exponential relaxation model (*gray traces*). The estimated relaxation time constants were between 47 and 123 ms (*blue numbers*). A faster component, indicated by the spikes in the residues (Fig 3B, *ϵ*(**L**_0_); Fig. 3C, *σ*_λ_(*ϵ*(**L**_0_)), *black trace*) is addressed in Supplemental discussion (Sec. 6.1).

### 3.3 Estimation of visual pigment absorption spectra

Spectral components **ū**_1..4_(**L**_C_) are shown in Figure 3D. The first spectral component **ū**_1_(**L**_C_) (Fig. 3D *blue trace*) was fitted with a visual pigment template model [^32^] for the absorption spectra of rhodopsin and metarhodopsin. Each template was parametrised with the position of the alpha peak *λ*_*α*_ and the relative magnitude of the beta peak *r*_*β*_. The alpha peak position implicitly defined the beta peak position and width (Sec. 7.6).

Component **ū**_1_(**L**_C_) was fitted with a model constrained with *r*_*β*M_ ≡ 0 (Fig. 3D, *gray trace*). The model estimated the alpha peak wavelengths of rhodopsin and metarhodopsin as **R547/M501** (nm), and the alpha peak absorbance ratio −*α*_M_*/α*_R_ = *q*_*α*_ ≈ 1.27. Omission of the beta peak of metarhodopsin (*r*_*β*M_ ≡ 0) was necessary because the shapes (i.e. the position and the bandwidth) of rhodopsin and metarhodopsin beta peaks are almost the same, and therefore hardly linearly separable *even if* the spectroscopic signal below 390 nm were acquired. The parameter estimate for rhodopsin beta peak relative magnitude is understood as absorbance *excess* of rhodopsin over the metarhodopsin beta peak (when positive), *r*_*β*Δ_ ≈ 0.19. Assuming exactly the same beta peak shapes, the two relative beta peak magnitudes can be related to the excess using *r*_*β*Δ_ = *r*_*β*R_ − *q*_*α*_ · *r*_*β*M_. Taking *r*_*β*R_ ≡ 0.25 and alpha peak ratio *q*_*α*_ = 1.27, the relative peak magnitude of metarhodopsin is *r*_*β*M_ ≈ 0.05.

The transient spectra {**a**} and the matched baseline spectra {**b**} are analogous to absorbance difference and sum, respectively. Using additional information from the baseline spectra could improve the parameter estimates, particularly for the beta peaks, which are otherwise inseparable due to having almost the same shapes. Three concatenations [**a**], [**b**], [**a, b**] were subjected to SVD and fitted with template models. The first two SVD spectral components of the concatenation of both types of spectra **ū**_1,2_([**a, b**]) contained about 66% and 24% of the signal, respectively (Fig. 4A, *blue, red trace*). ‘Temporal’ components 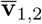 are shown separately for transient amplitudes and matched baselines (Fig. 4C, see *y-axis labels*). The contatenation was subjected to fitting with an unconstrained 4-parameter template model (Fig. 4A, *sky traces*), estimating **R552/M496** and *r*_*β*R_ ≈ 0.45, *r*_*β*M_ ≈ 0.31. The model constrained with *r*_*β*R_ ≡ 0.25 yielded **R552/M497** and *r*_*β*M_ ≈ 0.05 (Fig. 4A, *brown traces*). The residues of fitting the first component *ϵ*(**ū**_1_([**a, b**])) are comparable between the two models, while the residues of the second component *ϵ*(**ū**_2_([**a, b**])) are larger for the constrained model (*brown vs. sky trace, mauve arrow*). The first component of transient-only concatenation **ū**_1_([**a**]) contained about 89% of signal (Fig. 4B, *black trace*). The alpha peak estimates **R547/M501** obtained from fitting the parameter model with constraint *r*_*β*M_ ≡ 0 (*blue trace*) were used as constraints for fitting the dataset **u**_1,2_([**a, b**]), yielding beta peak estimates *r*_*β*R_ ≈ 0.44, *r*_*β*M_ ≈ 0.35.

**Figure 4.**
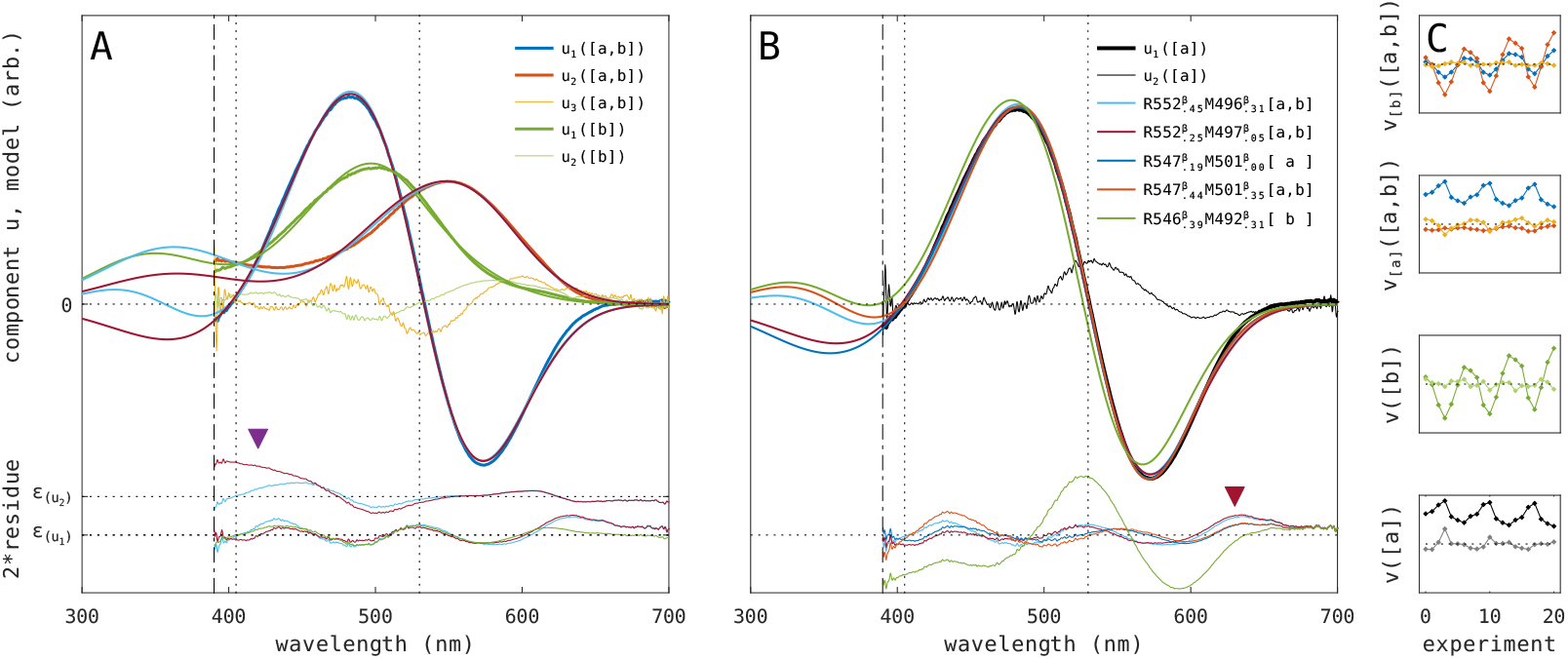
SVD components obtained from transient and baseline spectra, and template model comparison. **(A) Components of mixed provenience and of baseline provenience**. SVD components from concatenation **ū**_1,2,3_([**a, b**]) (*blue, red, yellow traces*). SVD components from baseline concatenation **ū**_1,2_([**b**]) (*green traces*). Residues *ϵ*_1,2_, *mauve arrow* points to **(B) Components of transient provenience**. Components **ū**_1,2_([**a**]) (*thick, thin black traces*). **Fitting residues** *ϵ* (AB, *thin traces, bottom*). *Mauve arrow* points to the wavelength range where the model with constraint *r*_*β*R_ ≡ 0.25 (*brown*) has larger residues than the unconstrained model (*sky*). *Brown arrow* points to the wavelength range where the models predict a higher than measured signal. **(C) Components** 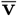. Dataset provenience on vertical axis label, axis span is equal in all four graphs. Component signals: *s*_[1:3]_*/*Σ*s* from concatenation of [*a*] : *{*0.887, 0.021, 0.014}, of [*b*] : {0.896, 0.025, 0.016}, of [*a, b*] : {0.663, 0.242, 0.014} **Models in (A**,**B)**: fit to **ū**_1,2_([**a, b**]) without constraints (*sky trace*); fit to **ū**_1,2_([**a, b**]) with constraint *r*_*β*R_ ≡ 0.25 (*brown trace*); fit to **ū**_1_([**b**]) with constraint *r*_*β*R_ ≡ 0.31 (*green trace*). **Models in (B)**: fit to **ū**_1_([**a**]) with constraint *r*_*β*M_ ≡ 0 (*blue trace*)); fit to **ū**_1,2_([**a, b**]) with alpha peak constraints obtained from the previous model (*red trace*). **Parameter estimates** (B, *legend*).

The predictions of the models fitted to **u**_1,2_([**a, b**]) were matched in the least squares sense to the SVD transient component **u**_1_([**a**]) in Fig. 4B (*red, sky, brown traces*). These models match the data better than the model R549/M492 obtained from fitting SVD of baseline-only concatenation **u**_1_([**b**]), as shown by a larger residue trace *ϵ*(**ū**_1_([**a**])) (Fig. 4B, *bottom green trace*). Due to the limited spectroscopic data range, a definitive model selection was not warranted, however, the unconstrained model **R552/M496** with *r*_*β*R_ ≈ 0.45, *r*_*β*M_ ≈ 0.31 (*sky trace*) seems to be the more plausible than the model with constraint *r*_*β*R_ ≡ 0.25, as it deviates less from the spectroscopic data below 450 nm (Fig. 4A, *ϵ*(**ū**_2_), *mauve arrow*). In the red wavelength range 600–650 nm, all models predict larger than measured transient signals (Fig. 4B, *ϵ*(**ū**_1_), *brown arrow*), possibly due to waveguide effects (see Sec. 6.2 & 6.3).

### 3.4 Statistics of template parameters

While the asymptotic errors of template parameters could be obtained from any non-linear fitting procedure on SVD component spectra, fitting individually the transient spectra and the pairs of transient and baseline spectra provided additional insights into the relation between the spectroscopic signal and the statistics of parameter estimates. The individual transient spectra {**a**} were fitted to the model constrained by *r*_*β*M_ ≡ 0. The parameter estimates for alpha peaks are shown along with the secondary estimates for the isosbestic and extrema wavelengths in Fig. 5A (*brown*). The alpha peak estimates were {*λ*_*α*R_ ≈ 546.6(1.5) nm, *λ*_*α*M_ ≈ 500.7(0.9) nm} (parenthesis contain standard deviations). The spectroscopic signal strength on the ordinate is represented by the transient amplitude (−*α*_R_) (Sec. 3.5). The estimates do not show an increased spread in the lower part of the tested signal range. Beta peak absorbance excess was estimated to *r*_*β*Δ_ ≈ .194(.023) (Fig. 5B, *brown*), expectedly similar to the estimates in Fig. 3D and Fig. 4B, as spectroscopic information and the model were essentially the same.

**Figure 5.**
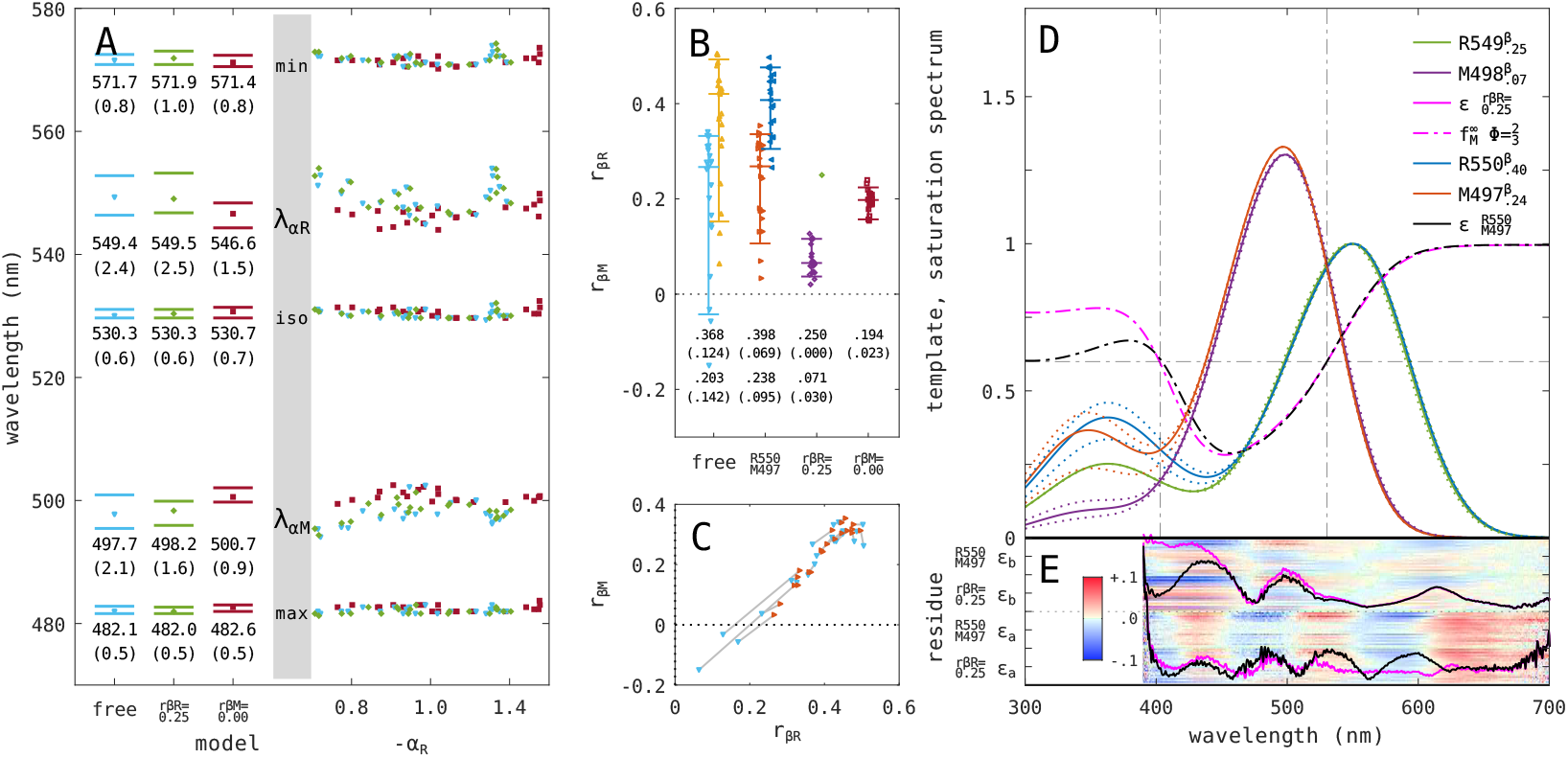
Statistics of template parameters. **(A) Characteristic wavelengths**. The wavelengths estimates from fitting: transient amplitudes {**a**}, model with constraint *r*_*β*M_ ≡ 0 (*brown squares*); pairs {**a, b**}, model without constraints (*sky triangles*); pairs {**a, b**}, model with constraint *r*_*β*R_ ≡ 0.25 *(green diamonds)*. Linear component −*α*_**R**_ is used as a proxy for the spectroscopic signal strength. Absorbance difference minimum (*min*), rhodopsin peak wavelength (*λ*_*α***R**_), main isosbestic point (*iso*), metarhodopsin peak wavelength (*λ*_*α***M**_), absorbance difference maximum (*max*). Quantiles [0.1, 0.9] (*bars*). *Numbers* show mean (standard deviation). **(B) Beta peak parameters**. The metarhodopsin and rhodopsin relative beta peak magnitude estimates *r*_*β*M_, *r*_*β*R_ from fitting the 4-parameter model on pairs {**a, b**} with the unconstrained model (*free: sky, yellow triangles*); the model with constraint R550/M497 (*red, blue triangles*); the model with constraint *r*_*β*R_ ≡ 0.25 (*mauve, green diamonds*). The rhodopsin beta peak absorbance excess estimate *r*_*β*Δ_ from fitting the model with constraint *r*_*βM*_ ≡ 0 on transient amplitudes {**a**} (*brown squares*). Quantiles [0.1, 0.9] (*bars*). *Numbers* show mean (standard deviation). **(C) Correlation of beta peak parameter estimates**. Estimates *r*_*β*M_, *r*_*β*R_ from the unconstrained model (*sky down triangles*) and the model with R550/M497 constraint (*red right triangles*). **(D) Visual pigment templates and predicted saturation spectra**. Templates from fitting pairs {**a, b**} with the model with constraint *r*_*β*R_ = 0.25 (rhodopsin: *green*, metarhodopsin: *mauve*); with constraint R550/M497 (rhodopsin: *blue*, metarhodopsin: *red*). Intervals shown as median (*solid traces*) and quartiles (*dotted traces*). Predicted saturation spectra (unconstrained model: *black dash-dotted trace*; constrained model: *pink dash-dotted trace*), using relative quantum efficiency Φ = γ_M_*/γ*_R_ = 2/3. Measured isosbestic wavelengths and predicted isosbestic saturation fraction 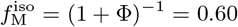 (*dash-dotted lines*). **(E) Fitting residues**. Residue standard deviation *σ*_*n*_(*ϵ*(**a**)), *σ*_*n*_(*ϵ*(**b**)): model with constraint *r*_*β*R_ ≡ 0.25 (*pink traces*); model with constraint R550/M497 (*black traces*, arb. scaling). *Heatmap*: fitting residues of individual transient and baseline spectra.

Next, the unconstrained model was fitted to the *pairs* of transient and matched baseline spectra {**a, b**}. The estimates for alpha peak wavelengths {*λ*_*α*R_ ≈ 549.4(2.4) nm, *λ*_*α*M_ ≈ 497.7(2.1) nm} moved slightly apart (Fig. 5A *sky*), compared to the model fitted to the set of transient spectra {**a**}. The two estimates for beta peak relative magnitudes had large statistical errors (Fig. 5B, *r*_*β*R_ *yellow, r*_*β*M_ *sky*). By constraining the alpha peak wavelength parameters, the beta peak parameters became more reliable, *r*_*β*R_ ≈ 0.37(0.08): *blue*; *r*_*β*M_ ≈ 0.22(0.12) (*red*). The scatter of *r*_*β*R_, *r*_*β*M_ estimated by these two models is shown in Fig. 5C. Finally, the model with the constraint *r*_*β*R_ ≡ 0.25 estimated the metarhodopsin beta peak relative magnitude as *r*_*β*M_ ≈ 0.07(0.03) (Fig. 5B, *mauve*).

Fig. 5D shows the predicted absorption spectra estimated from the two constrained four parameter fits to the pairs {**a, b**}. Both models predict well the experimentally observed isosbestic points at 405 and 530 nm (*vertical lines*). The two rhodopsin templates start differing below 460 nm, the two metarhodopsin templates below 440 nm. The fitting residues for the transient spectra and their ensemble standard deviations are comparable between the two models (Fig. 5E *ϵ*(**a**): *heatmap*; *σ*_*n*_(*ϵ*(**a**)): *pink, black traces*). Below 440 nm, the ensemble standard deviation of the baseline spectra *σ*_*n*_(*ϵ*(**b**)) is markedly larger for the model with beta rhodopsin peak constraint *r*_*β*R_ ≡ 0.25 (*ϵ*(**b**): *pink trace*) than for the model with constrained alpha peak wavelengths (*black trace*).

### 3.5 White light creates a high metarhodopsin fraction

Fitting the visual pigment template models to the pairs of spectra {**a, b**} yielded four linear component amplitudes per pair of spectra, (*α*_R_, *α*_M_, *β*_R_, *β*_M_). Components (−*α*_R_, *α*_M_) are shown in Fig. 6A (*red*). The linear slope laid over the data points yielded the metarhodopsin/rhodopsin alpha peak absorption ratio *q*_*α*_ ≈ 1.33(0.02). The ratio obtained from fitting the transient spectra {**a**} with the model constrained to *r*_*β*M_ ≡ 0 was *q*_*α*_ ≈ 1.27(0.02) (*black*).

**Figure 6.**
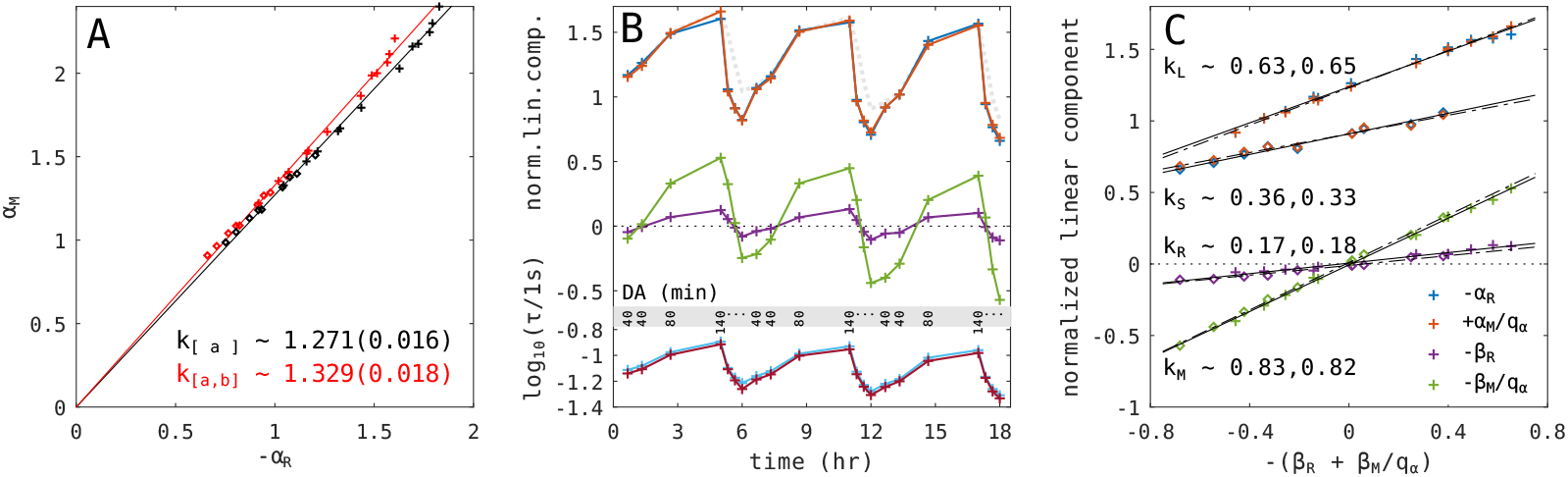
Analysis of linear components. **(A) Metarhodopsin to rhodopsin alpha peak ratio**. The slope of the relation between the rhodopsin (−*α*_R_) and the metarhodopsin component (*α*_M_) estimates the alpha peak ratio (*red*, combined dataset {**a, b***}* : *q*_*α*R_ ≈ 1.33 *±* 0.02) and (*black*, transient only dataset {**a**} : *q*_*α*R_ ≈ 1.27 *±* 0.02) **(B) Time course of the normalised linear components**. The components were obtained from the combined dataset {**a, b**}. The transient components of rhodopsin (−*α*_R_, *blue*) and metarhodopsin (*α*_M_*/q*_*α*_, *red*), the baseline components of rhodopsin (−*β*_R_, *mauve*), metarhodopsin (−*β*_M_*/q*_*α*_, *green*). Logarithm of the relaxation time constants of rhodopsin (log_10_ *τ*_R_, *sky*) and metarhodopsin (log_10_ *τ*_M_, *brown*). Preceding dark adaptation intervals (*black numbers*, minutes; *dots* 20 min). **(C) Relation between the normalised linear components and the total visual pigment content**. The four normalised linear components shown as a function of the proxy for the total visual pigment content −(*β*_R_ + *β*_M_*/q*_*α*_). Transient component of rhodopsin (*blue*), metarhodopsin (*red*), baseline component of rhodopsin (*mauve*), metarhodopsin (*green*). The dataset is divided into long (*crosses*) and short (20 min) preceding dark adaptation intervals (*diamonds*). The linear regressions (*black lines*) and their slopes (*black numbers*). Slopes of normalised transient components after long DA (*k*_L,M_ ≈ 0.63, *k*_L,R_ ≈ 0.65), short DA (*k*_S,M_ ≈ 0.36, *k*_S,R_ ≈ 0.33). Slopes for normalised matched baseline component for rhodopsin (*k*_R,L_ ≈ 0.17, *k*_R,S_ ≈ 0.18), for metarhodopsin (*k*_M,L_ ≈ 0.83, *k*_M,S_ ≈ 0.82). The rhodopsin and metarhodopsin baseline slopes sum to one as per definition of x-axis and correspond to post-exposure equilibrium fractions, 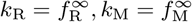.

The changes of the four normalised linear components over 18 hours are shown in Fig. 6B using component amplitude normalisation with *q*_*α*_ = 1.33 (Sec. 7.7). The normalised transient components of rhodopsin (−*α*_R_, *blue crosses*) and metarhodopsin (*α*_M_*/q*_*α*_, *red crosses*) were reaching a plateau in the exposures following long preceding dark periods. The metarhodopsin baseline component (−*β*_M_*/q*_*α*_, *green trace*) had a larger oscillation than its rhodopsin counterpart (−*β*_R_, *mauve trace*), indicating the prevalence of metarhodopsin in the post-exposure equilibrium. Relaxation rates were faster after short dark periods (6B, *brown and sky crosses*). Relaxation kinetics is further addressed in Sec. 6.1.

The normalised components are plotted in relation to −(*β*_R_ + *β*_M_*/q*_*α*_), the proxy for the total visual pigment content in the post-exposure equilibrium (Sec. 7.7). The transient components (Fig. 6C; rhodopsin: *blue*; metarhodopsin: *red*) are larger for long preceding dark intervals (40-140 min, *crosses*) than for the short intervals (20 min, *diamonds*). The equilibrium metarhodopsin fraction is estimated by the slope 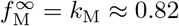 (*green symbols*) and is independent of the preceding dark interval.

For the exposures following long (L) dark intervals, when the rhabdoms presumably contain only rhodopsin, the slopes of normalised transient components *k*_L_ (*blue and red crosses*) should equal the estimated *k*_M_ ≈ 0.82. The discrepancy (*k*_L_ ≈ 0.64) was due to the experimental necessity of throwing away the first few spectra that were acquired during shutter transition time. Assuming that after long dark intervals, the pre-exposure rhodopsin fraction is indeed 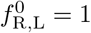, the transient amplitudes should be corrected by a factor *c* = *k*_L_*/k*_M_ ≈ 1.30, estimating the pre-exposure rhodopsin fraction for short (20 min) dark intervals to 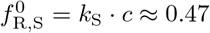. Increasing the rhodopsin fraction from 0.18 to 0.47 in 20 minutes and to one in 40 minutes is in line with the kinetics reported at room temperature [^87^].

Extrapolation of the slopes *k*_L_ estimated the axis intercept at {−(*β*_R_ + *β*_M_*/q*_*α*_), −*α*_R_} ≈ {−2, 0}. Using this intercept (*not shown*), the floating visual pigment content [−0.7 · · · 0.7] can be referenced to the range ≈ [1.3 · · · 2.7], expressed in terms of double-pass absorbance at the alpha peak wavelength of rhodopsin.

## 4 Discussion

A comprehensive analysis pipeline for the analysis of visual pigment photorelaxations was tested on the eyeshine reflectance spectra acquired using a spectrometer and broadband probing light from a living butterfly. The main (LW) bistable visual pigment of *Fabriciana adippe* was characterised by analysing 21 short exposures interleaved with dark adaptation intervals. Approximately ten seconds of UV-VIS spectroscopic signal were analysed. For each log-reflectance array, the exponential relaxation model extracted a transient spectrum and a baseline spectrum. The former describes photorelaxations during the exposures, the latter follows long-term changes of visual pigment content due to bleaching. The two types of spectra were subjected to singular value decomposition and template fitting, yielding rhodopsin and metarhodopsin absorption spectra.

### 4.1 Additional information in baseline spectra

Baseline matching procedure (Sec. 7.5) removed experimental effects (light source, microscope transmission, spectrometer sensitivity, corneal and tapetal transmittance, preparation drift), but kept spectral information related to the visual pigment concentration (Fig. 2). The visual pigment content components obtained from the matched spectra were following the long-term changes brought about by the photoreceptor’s machinery for pigment synthesis and removal (Fig. 6B). Matched baseline spectra crucially provided additional spectroscopic information that improved the parameter estimates for the beta peaks (Fig. 5B). Baseline spectra were to our knowledge not used in this manner in previous studies.

The importance of the additional information gained from analysing transient and matched baseline spectra together can be envisioned in the extreme case where the rhodopsin and metarhodopsin absorption spectra are exactly the same: the transient spectra would be zero, while the matched baseline spectra could still be fitted with a (single) template model. A showcase of linear inseparability, where the two pigment forms have virtually the same spectral shapes, is the visual pigment JSR1 of the jumping spider *Hasarius* [^88^]. Inclusion of matched baselines is expected to improve the estimation of alpha peak wavelengths and alpha peak absorption ratio in the species where the alpha peaks are close together.

Baseline rhodopsin and metarhodopsin linear components obtained from template fitting of the (matched) baseline spectra were unreferenced due to subtracting the ensemble average (Fig. 6C). By reëstablishing the zero reference, the total pigment content could be estimated to [1.3 · · · 2.6], expressed in terms of double-pass absorbance of rhodopsin. This zero-referencing procedure is more generally applicable than attempting to achieve a complete bleach as the reference [^10^]. The procedure should however be used with caution when the x-axis intercept is estimated from a limited data range, particularly when the data has higher scatter than in the case analysed here.

### 4.2 SVD facilitates visual pigment template fitting

Singular value decomposition (SVD) is a dimensionality-reducing matrix operation that has been employed in spectroscopy of vertebrate visual pigments [^85^] and cytochromes in living white-eyed blowflies [^95^]. SVD is often used in multivariate spectral analysis pipelines [^38–40,43,62^].

Here, SVD was first used to condition the reflectance spectra prior to the logarithmic transformation (Sec. 7.3). SVD was then applied to the large concatenated array of baseline-corrected transients (Fig. 3), and to the concatenations of transient and matched baseline spectra (Fig. 4). Between 80 to 90% of signal energy was contained in the components subjected to template fitting. SVD components produce less noisy spectra compared to plain arithmetic averaging, but at the peril of rejecting a relevant spectroscopic signal. SVD-based conditioning and component extraction are expected to play an important role in the analysis of narrow-band eyeshine reflections from butterfly eyes where ommatidia contain red filtering pigments that severely reduce the short-wavelength reflected light [^58,75,84^].

### 4.3 How precise are the estimated spectra?

The alpha peak wavelength estimates obtained through fitting data of different proveniences were up to 5 nm apart (**R547–551, M497–501**), while the standard deviations of estimates were up to 2.5 nm (Figs. 3, 4, 5). Some possible sources of estimate bias are listed in Supplemental Discussion (Sec. 6.2). Accuracy of the analysis pipeline should be assessed by comparing results from different eye regions of the same animal, and from different animals of the same species.

The beta to alpha peak magnitude ratio estimated from mixed provenience data (transient and baseline spectra) had relatively large errors (Fig. 5). This is to a large extent due to the lack of spectroscopic signal below 390 nm. The estimates were *r*_*β*R_ ≈ 0.40(0.07), *r*_*β*M_ ≈ 0.24(0.10). Constraining rhodopsin ratio to the standard value *r*_*β*R_ ≡ 0.25, the ratio for metarhodopsin was estimated to *r*_*β*M_ ≈ 0.07(0.03). Both sets of estimates are consistent with the measured isosbestic wavelength around 405 nm (Fig. 5). A similar isosbestic wavelength was indicated in the comma butterfly *Polygonia* [^87^], but not in the difference spectra of A1 visual pigments of fireflies measured in slices [^20^]. Intracellular measurements of green photoreceptors in butterflies reported that the relative beta peak magnitudes of rhodopsin in silver-washed fritillary *Argynnis paphia* and the black tiger *A. sagana* are between 0.3 and 0.7 [^42^], in the lesser purple emperor *Apatura ilia* 0.5, and in the two-tailed pasha *Charaxes jasius* 0.4 [^58^], but these values may not accurately represent true rhodopsin absorption spectra due to (self) screening or synaptic interactions.

The estimated metarhodopsin to rhodopsin alpha peak absorption ratio of *Fabriciana* was slightly lower *q*_*α*_ ≈ 1.27(0.02) for the transient dataset than for the dataset of mixed provenience, *q*_*α*_ ≈ 1.33(0.02) (Fig. 5AD). Table 1 lists some characterised bistable pigments. The reported alpha peak absorption ratio covers a large range, *q*_*α*_ = [1.2 · · · 2.0], possibly due to different measurement methods.

**Table 1.** Some invertebrate visual pigments. **Abbreviations**: *s*uperposition, *a*pposition, *c*amera eye optics, *dors*al; *chr*omophore: A3 hydroxyretinal, A1 retinal; *q*_*α*_ metarhodopsin to rhodopsin alpha peak absorption ratio; *meth*od: eye*shine, slice, extr*act, *sev*eral, *crys*tal; error (.00) indicates constrained models. **References**: G75^29^, C82^19^, M83^53^,S99^64^, C00^20^, V05^87^, S10^77^, M15^52^,V19^88^, P22^58^.

| | chr | genus | name<br>gene | $\lambda_{\alpha R}$<br>[nm] | $\lambda_{\alpha M}$<br>[nm] | $\frac{\lambda_{\alpha R}}{\lambda_{\alpha M}}$ | $q_\alpha$<br>(M/R) | meth | ref |
| --- | --- | --- | --- | --- | --- | --- | --- | --- | --- |
| a | A3 | *Fabriciana* |  | 550(5) | 500(5) | 1.10 | 1.30(.05) | shine |  |
| a | A3 | *Charaxes* | pasha | 543(6) | 502(3) | 1.08 | 1.27(.04) | shine | P22 |
| a | A3 | *Polygonia* | comma | 532 | 492 | 1.08 | 1.30(.00) | shine | V05 |
| a | A3 | *Apatura* | emperor | 526(3) | 496(4) | 1.06 | 1.25(.00) | shine | P22 |
| a | A3 | *Drosophila* | R8y rh6 | 515 | 468 | 1.10 | 1.2 | extr | S99 |
| s | A3 | *Deilephila* | hawkmoth | 520 | 480 | 1.08 | 1.7 | extr | H73 |
| s | A3 | *Galleria* | waxmoth | 510 | 484 | 1.05 | 1.43(.05) | slice | G75 |
| s | A1 | *Photuris* | firefly | 545 | 485 | 1.21 | 1.6(0.4) | slice | C00 |
| s | A1 | *Photinus* | firefly | 555 | 485 | 1.14 | 1.8(0.2) | slice | C00 |
| s | A1 | *Orconectes* | crayfish | 535 | 510 | 1.05 | 1.4 | slice | C82 |
| c | A1 | *Hasarius* | spider jsr1 | 535 | 535 | 1.00 | 1.3 | crys | V19 |
| c | A1 | *Todarodes* | squid | 480 | 480 | 1.00 | 1.3 | crys | M15 |
| c | A1 | *Eledone* | octopus | 470 | 516 | 0.91 | 1.5 | extr | H73 |
| a | A1 | *Apis* | drone dors. | 445 | 505 | 0.88 | 1.7 | slice | M84 |

Statistics of alpha peak parameter estimates is often not reported, neither in the spectroscopic studies in vivo [^9–11,64,87^], nor in the studies on cell expression systems, where the spectroscopic signal is more noisy [^47,93^]. It would be prudent if the spectroscopic parameters of rhodopsin *and* metarhodopsin forms of bistable pigments – also for their beta peaks – were reported with error intervals. A systematic study relating alpha and beta peak positions, peak extinction, quantum efficiency and oscillator strengths of bistable pigments is not known to have been performed.

### 4.4 Saturation spectrum can be predicted and measured

The acquired spectral signal did not extend below 390 nm, precluding accurate estimation of beta peak parameters. In this species, using a UV-extended illumination source and an epi-illumination objectives with good UV transmission might extend the signal to about 350 nm. Many butterfly species however do not reflect UV and blue light due to pigment and tissue absorption, or due to narrow reflection of the tapetum.

Adding an adjustable monochromatic light source would enable mapping the saturation spectra that predict the metarhodopsin fraction achieved by sufficiently long (saturating) exposure to monochromatic light. Saturation spectrum is defined as 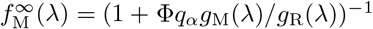, where *g*_M_(*λ*), *g*_R_(*λ*) are normalised metarhodopsin and rhodopsin absorption spectra, respectively, and the relative quantum efficiency Φ = *γ*_M_*/γ*_R_ is assumed to be wavelength-independent [^19,77,80^]. The transient amplitude components estimated from the exposures following saturating preadaptations could be used to construct the saturation spectrum; alternatively, the fraction of metarhodopsin could be measured through its fluorescence [^5,17,18,82^]).

Saturation spectra of two template pairs are shown in Fig. 5D. Using Φ = 2*/*3, the models with higher and lower beta peaks respectively predict a measurable difference in the saturating metarhodopsin fraction in the UV range. The minimal fraction of metarhodopsin *f*_M_ ≈ 0.3 is predicted to follow saturation with blue light around 450 nm.

### 4.5 Method improvement and comparison

Spectroscopic methods used for measuring absorption spectra of visual pigments in living butterflies can be broadly classified into those employing attenuated monochromatic light and a photon counter [^9–11^], and those employing a broadband source and a spectrometer [^58,87^]. Broadband spectroscopy approach used here can be implemented on a standard epi-illumination microscope, and can potentially be made field-portable. The method yields spectra with high signal to noise ratio, particularly when compared with heterologous cell expression systems. A clear advantage of the *in vivo* approach is the absence of wet lab procedure and a long experimental time window that enables employment of the fully functional visual pigment synthesis/removal cycle to broaden the range of experimental conditions. The main disadvantages are the spectroscopic range limited by screening pigments, tissue absorption and tapetum reflectance, and possible crosstalk from the other visual pigments.

The presented method can be applied to the majority of butterfly species from the families Nymphalidae, Lycaenidae and Riodinidae. In some species of these families, and in the species of the family Pieridae, the ommatidia in the ventral eye regions have red screening pigments [^42,75^] that may limit spectroscopic signal above the metarhodopsin peak, rendering template fitting less reliable. In large butterflies, optical density may be too high due to long rhabdoms. Intentional bleaching caused by saturating red light after each exposure may improve the spectroscopic signal of the LW visual pigment, and may unmask the UV and blue visual pigments [^9,87^]. On the other hand, applying saturating blue light after each exposure would minimize bleaching and enable using a faster exposure sequence. For example, using three minute dark intervals to allow pupil to open [^81^], ten seconds of spectroscopic data could be collected in about an hour. The method could be complemented with measurements of saturation spectra and with electrophysiological and optophysiological recordings of spectral sensitivities [^58^]. Modification of template fitting procedure to spectral sensitivity datasets is addressed in supplemental discussion (Sec. 6.3).

### 4.6 Application to other insect groups

The butterfly family Hesperidae (skippers) and many moth families exhibit a bright eye glow. In their superposition eye design, the distal optics is shared between the ommatidia, increasing the light flux to the rhabdoms, and the relaxation rate for up to hundred times [^7,94^]. The rate can be reduced using an optical system that limits the incoming light field to a few facet lenses, while keeping a large aperture. This can be achieved with the optical system used for ORG measurements [^57,75^].

Genetic tools are being developed for Lepidoptera [reviewed in ^16,48^], and transgenic lines have been created for a few butterflies from the family Nymphalidae and moths from the families Pyrallidae and Erebidae [^69,70^], but this is not yet a routine endeavour; a few years more shall pass before the power of butterfly eyeshine spectroscopy will be combined with transgenic manipulation of visual pigments.

Transgenic fruitflies have been shown to express visual pigments from butterflies and beetles. Their sensitivity spectra have been measured through pupil action [^27^] and extracellular electrophysiology (ERG) [^67,68^]. Broadband spectroscopy in intact fruitfly eyes would be a good complementary method to study natural and genetically modified bistable visual pigments. Flies have open rhabdom apposition eyes, with photoreceptors R1-6 forming peripheral rhabdomeres, and photoreceptors R7 & R8 being stacked in the central rhabdomere. Flies do not have a tapetum, but the light backscattered from white-eyed blowflies bears spectroscopic signal from respiratory *and* visual pigments [^95^]. The signal in white-eyed fruitflies is lower due to their smaller eyes^∗^, so concatenation of exposures (Fig. 3) might be helpful.

A superior approach applicable to flies is to use antidromic (back) illumination, achieved by focussing light onto the head capsule, or inserting a light fibre behind the eye, resulting in an image of the seven rhabdomeres superimposed from several ten ommatidia at the eye centre [^25,37^]. This approach has been used for absorption spectroscopy [^72^] and pulse-probe spectroscopy [^46,61^]. Many insects have dense dark pigments below the retina, precluding use of antidromic illumination, but saturation spectra might still be mappable through measuring metarhodopsin fluorescence, as in flies [^5^].

## 5 Summary and outlook

A comprehensive analysis pipeline for studying the photorelaxations of visual pigment spectra obtained from the eyeshine reflections in intact butterflies is presented. The analysis pipeline was tested on the spectroscopic data obtained from a female brush-foot butterfly, high brown fritillary *Fabriciana adippe*. The main visual pigment was estimated through the templates of Govardovskii as rhodopsin **R547–R552** and metarhodopsin **M496–M501**, with the metarhodopsin to rhodopsin alpha peak absorption ratio 1.25–1.35. The transient spectra had isosbestic wavelengths at about 530 and 405 nm, the latter wavelength indicating that the rhodopsin absorption in the UV is higher than that of metarhodopsin. The relative magnitude of rhodopsin beta peak seems to be higher than the often assumed 0.25, and is provisionally estimated to about 0.30–0.45 for rhodopsin and 0.10–0.30 for metarhodopsin. The (single-pass) absorbance of rhodopsin alpha peak was estimated to about 1.3 in fully dark-adapted state.

Transient (absorbance difference) and baseline (absorbance sum) components of photochemical relaxations were for the first time analysed together, improving the separability of rhodopsin and metarhodopsin spectra. Statistical errors of the beta peak estimates could be further reduced by using a light source with the spectrum extending into the UV. The spectroscopic method could be expanded using preadaptation with saturating monochromatic light, and complemented with optophysiologically and electrophysiologically measured rhodopsin spectral sensitivity data that could be analysed with the same global fitting procedure. The estimated visual pigment templates could be compared against the opsin sequences in genomic databases. Butterfly eyeshine might be explored as a preparation for in-vivo pulse-probe spectroscopy to study the intermediate states of visual pigments. The optical setup could be adapted to other insects, particularly to white-eyed fruitflies expressing transgenic opsins, expanding the possibilities for studying molecular correlates of visual pigment tuning.

## Acknowledgements

Kentaro Arikawa for the sbfSEM picture; Gary D. Barnard, Gregor Belušič, Anna-Lee Jessop, Andrej Meglič, Michael W. Perry, Doekele G. Stavenga and Bodo D. Wilts for their suggestions and corrections.

## Supplemental material

## 6 Supplemental results and discussion

### 6.1 Complex relaxation kinetics in optically dense rhabdoms

The baseline-corrected transient arrays {**L**} were individually fitted with the template model with constraint *r*_*βM*_ ≡ 0 (Sec. 7.6) in order to yield independent estimates for rhodopsin and metarhodopsin relaxations. Seven pairs of relaxation traces α_**R**_(*t*), α_**M**_(*t*) are shown in Fig. 7A (*pairs in the same colour*). The relaxation traces of rhodopsin and metarhodopsin were fitted with the first order exponential relaxation model (Sec. 7.4). The relaxations were slower in the exposures following long dark intervals (Fig. 6B, log_10_ *τ*_R_, *sky*, log_10_ *τ*_M_, *brown*). The dataset was divided into the exposures following short (20 min) dark intervals (*diamonds*) and long dark intervals (*crosses*). The relation between the pairs of time constant estimates is shown in Fig. 7B. Metarhodopsin apparently relaxed about 5% faster than rhodopsin. A simple two-state rhodopsin-metarhodopsin photochemical model predicts the rates to be identical. The discrepancy could be due to relaxations of blue or UV visual pigments in photoreceptors R1 & R2 [^87^]. The relaxations got faster as the pigment content decreased, but the recorded data range is too narrow to predict the asymptotic relaxation rate in the low pigment content limit. This could be easily overcome by implementing intentional bleaching with saturating red light (Sec. 4.5).

**Figure 7.**
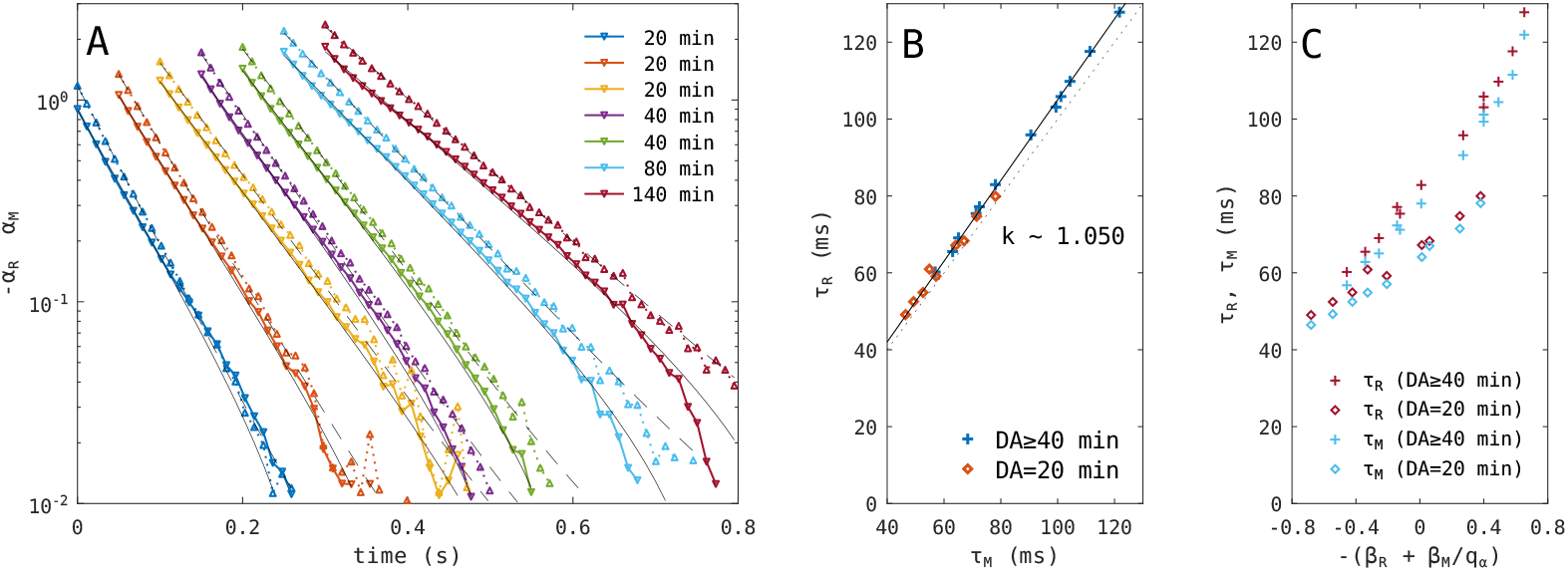
Relaxation time constants. **(A)** Semi-logarithmic plot of timecourses of rhodopsin components −*α*_R_(*t*) (*down triangles*) and metarhodopsin components *α*_M_(*t*) (*up triangles*) obtained from fitting seven baseline-corrected transient arrays {**L**} with a 3-parameter template model. Exponential relaxations fitted to the component timecourses (*black solid & dashed traces*). The modelled traces are not straight because the estimated relaxation equilibria were not constrained to zero. The timecourses are laterally displaced by 50 ms. **(B)** Relation between the pairs of relaxation time constants. Rhodopsin time constants are about 5 % slower than those of metarhodopsin. **(C)** Relation between the estimated visual pigment content and the time constants (rhodopsin, *brown*; metarhodopsin, *sky*). Exposures after short DA (20 min, *diamonds*) and long DA (40-140 min, *crosses*).

Relation between the relaxation time constants and the proxy for the total visual pigment content −(*β*_R_ + *β*_R_*/q*_*α*_) is shown in Fig. 7C (*τ*_*R*_ *sky, τ*_*M*_ *brown*). At the same pigment content, the relaxations following short and long dark intervals were respectively faster and slower (Fig. 7C). One might expect the opposite. Butterfly rhabdom is an optically dense medium, where the relaxation rate is falling along the rhabdom depth. After shorter dark intervals, the rhabdoms have a high fraction of more absorbing metarhodopsin, so the relaxation rate in the rhabdom depth should be lower, resulting in an overall reduction of the *apparent* (average) relaxation rate. On the other hand, the proxy for the total visual pigment content −(*β*_R_ + *β*_M_*/q*_*α*_) might be overestimated for the relaxations following short dark intervals, resulting in a shift of the points to the right (Fig. 7C, *diamonds*);

A fast relaxing component with time constant below 20 ms was indicated in the sharp temporal peaks in the residues of the initial relaxation model (Fig 3B, residues *ϵ*(**L**_0_); Fig. 3C, *σ*_*λ*_(*ϵ*(**L**_0_)), *black trace*). The fast relaxing component could be due to the intermediate **N** on the metarhodopsin to rhodopsin pathway that in the blowfly thermally decays with lifetime 13 ms [^33,61,73^]. The fast component did not resolve into a separate SVD component, but remained as a crosstalk in the first temporal SVD component (Fig. 3C, 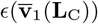, *pink trace*). The fast component was deemed too small to significantly influence the baseline estimation. If this was not the case, the relaxation model could be fitted to front-truncated spectra {**L**_0_}. A relaxation model with two components was tested, but it was not possible to reliable resolve the two time constants (*not shown*). In the experiment, this could be overcome by minimizing the shutter transition time and increasing the acquisition rate. In the analysis, advanced multivariate curve resolution techniques should be used [^43,62^].

Rhabdoms are complex optical media due to their length and optical density, the presence of additional UV- and blue-peaking rhodopsins in photoreceptors R1 and R2, and the waveguide effects. Theoretical analysis of relaxation kinetics in optically dense media have been conducted for chemical photosystems [^90,91^] and for vertebrate photoreceptors [^34^]. Understanding the complex interplay of relaxation rates and pigment content along the rhabdom depth would require additional experimentation and thorough modelling. Computational models that have calculated rhodopsin absorption along the rhabdom length in order to predict spectral sensitivities have accounted for either waveguide mode effects [^78^] or for metarhodopsin absorption [^6,57^], but not for both effects together.

### 6.2 Sources of pigment template parameter errors

The transient and baseline spectra and the SVD components obtained from them were virtually free of noise, but the template parameters estimated from these spectra might still be wrong. Some of the possible sources of parameter bias are listed here.

1. The exposures after long dark intervals produced noisier baseline spectra in the range 450 to 500 nm (Fig. 2). The log reflectance in equilibrium was low in this range due to the abundance of created metarhodopsin. As a single dark reference measurement is used, an error in it may cause a bias in the spectra acquired throughout the experiment (Sec. 2, Fig. 8B, distance between *green* and *blue* trace). The noise in the baseline spectra is expected to be higher in the species with large eyes, but could be overcome with intentional bleaching using red light (see Sec. 4.5).
2. If the photoreceptors R3-8 contained two visual pigments [^3,12,22,50^], this would intuitively result in the predicted alpha peak wavelengths to be between the two true values. If the two visual pigments had different relaxation rates, this would result in temporally unstable isosbestic points, similarly as reported for the crosstalk from photoreceptors R1 & R2 [^87^].
3. The measured reflectance spectra are a sum of reflectances from several ommatidia. Already within a single ommatidium, the relaxation rate and equilibrium fraction are functions of rhabdom depth due to the changing light spectrum, so a single exponential is not an accurate relaxation model. This can lead to inaccurate template estimation, particularly in the species with a complex eyeshine where some ommatidia have a red screening pigment [^6,42,58^] and the apparent relaxation rates between the ommatidia are expected to be markedly different.
4. About 20% of spectroscopic signal in the higher SVD components was excluded from template fitting, while the included spectroscopic signal, e.g. transient and baseline spectra, could be weighted differently. These arbitrary choices may influence the estimated template parameters.
5. The measured absorbance difference spectra may be modified by waveguide effects [^78^], resulting in a smaller absorbance difference in the red wavelength range (Fig. 4). This may cause a bias in the estimator for the rhodopsin peak wavelength. The issue is further addressed in Sec. 6.3.
6. The pigment templates of Govardovskii might not accurately describe the absorbance spectra of A3-based visual pigments. The templates have been developed for A1 and A2 rhodopsins [^28,32,83^]. It is assumed that the A1 rhodopsin template is valid for both A3 rhodopsin *and* A3 metarhodopsin. A correction for A3 metarhodopsin of fly Rh1 was given in [^77^]. The presented method can yield spectroscopic data across many butterfly species, and might enable a new parametrisation of A3 rhodopsin and metarhodopsin templates.

**Figure 8.**
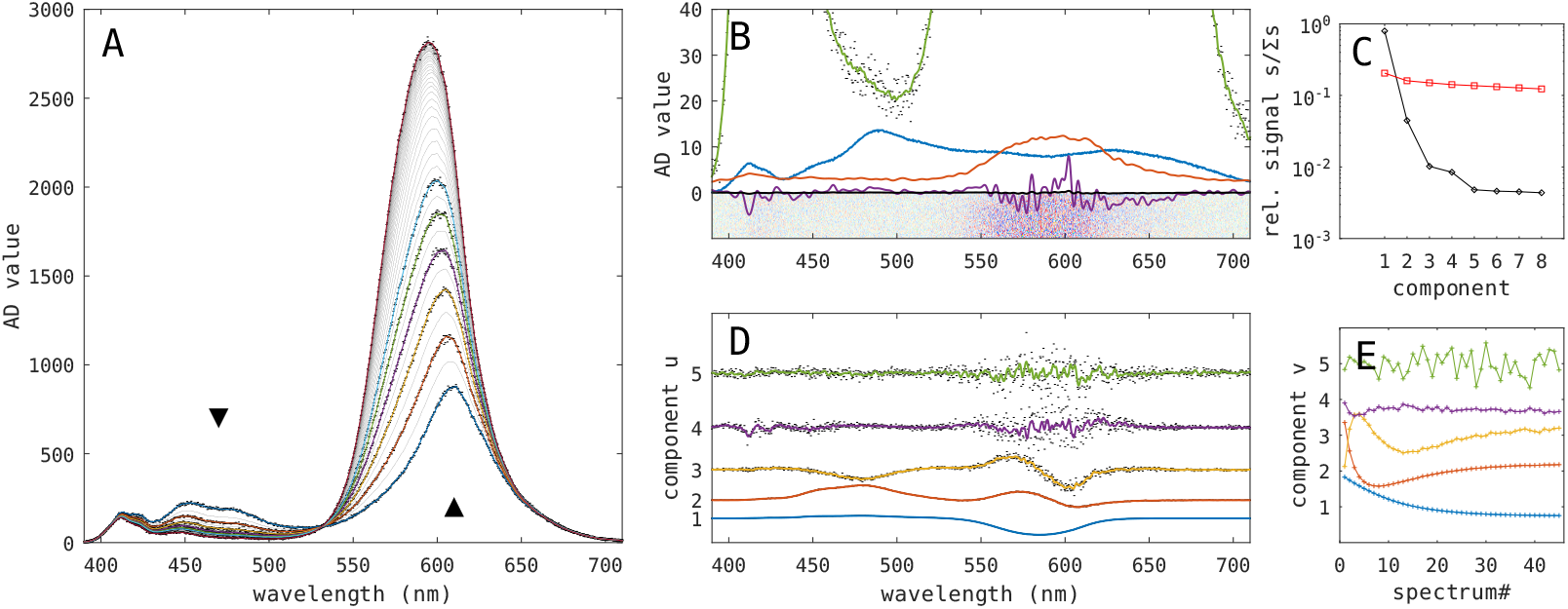
Conditioning of reflectance spectra. **(A)** The spectra in the conditioned array 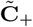 (*gray and coloured traces*). The first few and the last spectrum from the unconditioned array **C** (*black dots*). The minimal signal at 500 nm is around ADC value 20. **(B)** Residuals 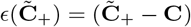 (*heatmap*). Low-pass filtered temporal mean of the residual 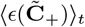 (*black trace*; 20*×* scaled, *mauve trace*), standard deviation 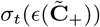 (*red trace*). A conditioned spectrum (*green trace*) laid over an unconditioned spectrum (*noisy black points*). Dark reference spectrum (*blue trace*). **(C)** Relative signal (*s*_*k*_/Σ*s*)(**M**) in the first few SVD components (*black diamonds*), signal energy remaining in higher components (Σ_[(*k*+1):]_*s/*Σ*s*)(**M**) (*red squares*). **(D)** Unscaled spectral components **u**_1..5_(**M**), showing shot noise in the higher components. **(E)** Unscaled temporal components **v**_1..5_(**M**).

### 6.3 Amendments to the global fitting procedure

In this study, the reflectance spectra were log-transformed before fitting relaxations and pigment templates, presumably removing all static non-linearity of the raw data. The design matrix used in the template fitting procedure (Sec. 7.6) therefore contained only two vectors with the normalised templates of rhodopsin and metarhodopsin, **X**_L_ = [**g**_R_, **g**_M_].

#### Non-linearities

To account for remaining non-linearities, the design matrix could include higher order and interaction terms of the normalized templates, e.g. 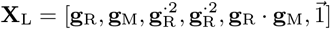, at the expense of noisier components returned in **B**_L_.

#### Waveguide effects

The measured transient spectra exhibited a smaller absorbance difference in the long wavelength range than what the templates predicted (Fig. 3D, 4B). The discrepancy might be due to waveguiding in the fused rhabdoms of butterflies. Only the fraction of light transported inside the rhabdom can be absorbed by the visual pigment. A model for the small white *Pieris* [^78^] predicts that the rhabdom with circular diameter 2 µm at 600 nm allows two waveguide modes with average fraction travelling inside 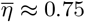, implying that the templated absorbance difference spectrum should be corrected to approach zero faster in the red range (Fig. 4B). In order to account for the reduced fraction of light amenable to absorption by the visual pigments due to waveguiding, the design matrix (Sec. 7.6) could be modified to 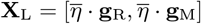, where the fraction *η*(*λ*) could be predicted from the waveguide theory [^36,76,78^], given as a heuristic function with fixed parameters [^2^], or modelled as a short-pass wavelength filter function with nonlinear parameters defining the transition band subjected to non-linear optimisation.

#### Spectral sensitivity datasets

Intracellular electrophysiological measurements of photoreceptor spectral sensitivity provide crucial information on the absorption spectra of rhodopsins [^6, 42^], albeit with added layers of complexity e.g. due to screening and synaptic interactions. The static response non-linearity is removed by recording a dose-response (log intensity-voltage) calibration and fitting it with a sigmoid function; the responses obtained from the isoquantal spectral runs are then transformed with the sigmoid inverse. Pupil response [^81^] is used in optical retinography (ORG) to measure the spectral sensitivities of several hundred individual ommatidia [^57,58^]. The pupils of photoreceptors are independent, so the compound pupillary sensitivity of each ommatidium can be approximated as a sum of spectral sensitivities of the contributing photoreceptors. Correction for pupil non-linearity is similar as for electrophysiological recordings. In the analysis of optophysiological and electrophysiological spectral sensitivity data, the negative linear component amplitudes can be rectified before calculating the residues inside the template fitting iteration loop. An additional (negative) metarhodopsin template would likely be needed to account for screening effects in electrophysiological spectral sensitivity datasets. The minimal design matrix for a dataset with UV, blue and green photoreceptors and a single metarhodopsin without synaptic interactions could be 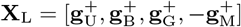, where the superscript plus signifies rectification.

## 7 Supplemental data analysis methods

### 7.1 Notation

Vectors are in bold, arrays and matrices in bold capitals, **a, L**. Transpose operation is indicated as **B**^*T*^ . Braces indicate sets {**a**}, brackets concatenation [**a, b**]. Fitting a set {**a**} and a concatenation [**a**] yields multiple sets and a single set of non-linear parameters, respectively. Ranges are indicated with a colon, e.g. **a**_[1 : 5]_. With the intent to help with coding, the array dimensions are indicated between corners: **A**^⌞*λ,t,n*⌝^ is an array with successive vertical, horizontal and page dimensions of wavelength, time and experiment, respectively. Fit residues are indicated as *ϵ*(**L**_0_). Average and standard deviation over a dimension are indicated with a subscripted chevron and sigma, ⟨**b**⟩_*λ*_ and *σ*_*n*_(*ϵ*(**L**)). Parameter estimates in the text are reported as averages with standard deviation in parenthesis, *m*(*σ*).

### 7.2 Singular value decomposition

Singular value decomposition (SVD) was then performed using function `[U,S,V]=svd(M,`’`econ`’) that decomposed a spectro-temporal matrix **M**^⌞*λ,t*⌝^ = **U**^⌞*λ,c*⌝^**S**^⌞*c,c*⌝^(**V**^⌞*t,c*⌝^)^*T*^ . The matrices **U, V** contained the spectral vectors **u**_[*c*]_ and temporal vectors **v**_[*c*]_, respectively. The diagonal of S contained scaling factors *s*_[*c*]_ sorted in a descending order. The temporal and spectral vectors are scaled as 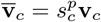 and 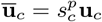. Using scaling (*p* = 1/2), recomposition simplifies to 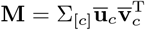. In Fig. 3 and 4 the components are shown with scaling (*p* = 1/2), in Fig. 8 without scaling (*p* = 0).

### 7.3 Signal conditioning

The acquired spectra contained raw analog-digital converter code/count values (ADC values) in the range 0-4095, with the ADC offset around 200. The spectro-temporal arrays first had the electronic dark and the optical dark references subtracted. The resulting arrays **C**^⌞*λ,t*⌝^ (the superscripted corners show the order of dimensions) contained 45 time points, covering 500 ms of the exposure after the shutter was fully open and before the pupil started closing, and 934 spectral pixels covering the wavelength range from 390 to 710 nm. The temporal average of the array was lowpass-filtered using a phase preserving low pass filter (functions `filtfilt, fir1(16,1/4)`), and subtracted from the original array, **M** = **C** − ⟨**C**⟩_*t*_.

The unscaled component spectra **u**(**M**), temporal components **v**(**M**) and scaling factors *s*(**M**) of a decomposed spectro-temporal matrix **M** are shown in Fig. 8CDE. The spectral components **u** were lowpass-filtered (`fir1(8,1/2)`) prior to recomposition. The temporal average was added back to the rank-reduced matrix recomposed from the first five SVD components, 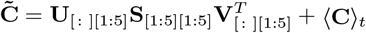.

The recomposed matrix was rectified (its elements made positive) by transforming the elements below threshold (*c < c*_*t*_) using function *c*_+_ = *c*_*f*_ + (*c*_*t*_ − *c*_*f*_)(1 + tanh(*c/c*_*t*_ − 1)), with the threshold and floor parameters *{c* = 3, *c* = 10^−3^} chosen by eye to correspond to the noise statistics of the raw spectra. The conditioned and rectified spectra in 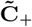 are shown in Fig. 8A (*coloured and gray traces*) along with a few unconditioned spectra in **C** (*black dots*). The procedure removed electronic noise over the whole range and shot noise between 550 and 600 nm (Fig. 8B, *red trace*), and did not introduce bias (*mauve trace*). The array was transformed to 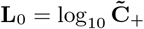. Its elements contained log-reflectance values analogous to absorbance measured in double-pass configuration. Rectification insured that the noise in the low signal parts of the spectra was not excessively amplified by the transformation. A few log-reflectance spectra from a single exposure are shown in Fig. 2B (*black traces*).

### 7.4 Global fitting of exponential relaxations

Exposure to light causes the bistable photopigment system to undergo relaxation described by 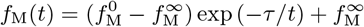, where the metarhodopsin fraction *f*_M_ relaxes from its starting value 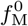 to its equilibrium value 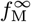 with a time constant *τ* proportional to the light intensity [^77^]. In the dark-adapted rhabdom, the starting fraction is *f* 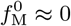, while the post-exposure equilibrium fraction 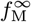 depends on the illuminant spectrum of the probing light. Despite the relaxation process with broadband illumination being more complex because both the relaxation rate and equilibrium metarhodopsin fraction change along the rhabdom, the measured (apparent) relaxation is quite accurately described by a simple exponential function [^87^]. The relaxations obtained in the measurements here are heuristically described by the equation *L*_0_(*λ, t*) = *a*(*λ*) exp(−*t/τ*) + *b*_0_(*λ*).

The global fitting procedure was implemented as an iterative optimisation performed over the temporal dimension of the log-reflectance spectra. The nested non-iterative ordinary matrix least squares (MLS) calculated by function `ols` returned linear components in the wavelength dimension. The MLS residues were in turn iteratively minimised in the outer loop through the optimisation of the non-linear parameters with the function `residmin`. The procedure returned the relaxation time constant as the non-linear parameter, and two spectra – the transient amplitude spectrum and the baseline spectrum. The log-transformed array **L**_0_ from a single exposure, containing 45 spectra, served as the input. The non-linear model described the relaxation in the temporal dimension. The input spectra were modelled by the linear equation 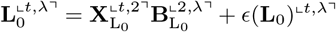 where 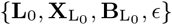 were the input, design, solution and residue matrices, respectively. The design matrix 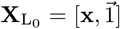 contained the vector **x** = exp(−**t***/*10^*τ*^) and a vector of ones. The timestamps **t** were read from the spectrometer acquisition data. The single non-linear parameter, the relaxation time constant, was parametrized as log_10_ *τ* for the purpose of numerical stability. The solution matrix returned two spectra, 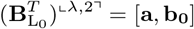. The transient amplitude spectrum **a** contained information on the (fast) photochemical relaxation, while the (raw) relaxation baseline spectrum **b**_0_ contained information on the visual pigment content in the equilibrium at the end of exposure. Each baseline-corrected transient array was obtained by subtracting its estimated baseline spectrum, **L** = **L**_0_ − **b**_0_. The residues *ϵ*(**L**_0_) are shown in Fig. 3A (*upper half*). The time courses contained in the baseline-corrected arrays **L** at four wavelengths are exemplified in Fig. 3B. Subtracting the single first or last spectrum from the array instead of subtracting **b**_0_ produced arrays with comparably higher noise (*not shown*). Exponential relaxations were also fitted to the timecourses of linear components *α*_R_(*t*), *α*_M_(*t*) obtained from fitting visual pigment templates (Sec. 7.6) to the arrays {**L**} (Fig. 7).

### 7.5 Matching of baseline spectra

The relaxation baseline spectra **b**_0_ obtained from fitting exponential relaxations (see 7.4) were similar above 620 nm (Fig. 2C). The small vertical offsets in this range were likely due to a slight drift of the preparation during the long hours of the experiment. The vertical offsets were removed by individually subtracting the average in the far red range λ_FR_ = [666..696 nm], 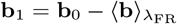. Each matched baseline spectrum was calculated by subtracting the ensemble average baseline spectrum, **b** = **b**_1_ − ⟨**b**_1_⟩_*n*_.

### 7.6 Global fitting of visual pigment templates

Template fitting procedure returned the iteratively optimised non-linear template parameters describing the templates for rhodopsin and metarhodopsin, and for each input spectrum [*j*], the linear component amplitudes corresponding to each template. The matrix equation defining the model was 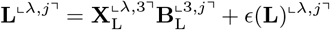. (Note the exchanged dimensions between 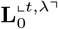 and **L**^⌞*λ,j*⌝^.) The design matrix contained vectors with the normalised A1 templates of Govardovskii for rhodopsin and metarhodopsin isoforms, and a vector of ones, 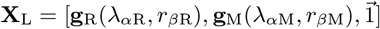. Each template was parametrised by two non-linear parameters: the position of the alpha peak *λ*_*α*_ and the relative magitude of the beta peak *r*_*β*_. The first parameter implicitly defined the position of the beta peak (around 350 nm) [^**32**^]. The output of the fitting procedure was a set of four non-linear parameter estimates *{λ*_*α*R_, r_*β*R_, *λ*_*α*M_, *r*_*β*M_}. In some instances, the constraints *r*_*β*M_ ≡ 0 or *r*_*β*R_ ≡ 0.25 were used, or the alpha peak positions *λ*_*α*R_, *λ*_*α*M_ were constrained to the values obtained from previous fits. The solution matrix **B**_L_ returned (temporal) vectors with linear component amplitudes of rhodopsin and metarhodopsin, and an offset vector, 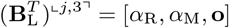. The offset was deemed to be negligible after initial trials and was omitted from the design matrix.

### 7.7 Normalisation of linear components

The spectral array that served as the input to the template fitting (see 7.6) could contain spectra of different provenience: those related to transients and *absorbance difference* (concatenated baseline-corrected spectral arrays **L**_C_, transient amplitude spectra **a**, or their SVD-derived spectral components), those related to baselines and *absorbance sum* (matched baselines **b** and their SVD-derived spectra), or those of mixed provenience, e.g. SVD components **ū** ([**a, b**]).

When the vectors in the input matrix were of transient provenience, the obtained linear component elements in the solution vectors were proportional to the *change* of pigment content during relaxation, 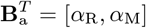. The changes were negative and positive, respectively, *α*_R_ *<* 0 *< α*_M_, indicating that rhodopsin content decreased and metarhodopsin content increased during the exposure.

When the spectral vectors in the input matrix were or baseline provenience, the obtained linear components were reporting floating (unreferenced) visual pigment content in the equilibrium after the exposure. Lower values of the linear component elements in the vectors of 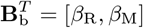 meant less reflected light and thus indicated a higher visual pigment content in the equilibrium.

The four linear components were normalized using *{*−*α*_R_, +*α*_M_*/q*_*α*_} and {−*β*_R_, −*β*_M_*/q*_*α*_}. The metarhodopsin content was corrected with the metarhodopsin to rhodopsin alpha peak absorption ratio *q*_*α*_ ≈ 1.3 estimated from the slope of the relation *k*(α_M_, −*α*_R_), see Fig. 6A. Due to the baseline matching procedure, the components *β* were without a zero reference point, the term −(*β*_*R*_ + *β*_*M*_ */q*_*α*_) served as a proxy for the (unreferenced) total visual pigment content in the equilibrium.

### 7.8 Steps in the analysis procedure

#### Signal conditioning

1. Reflectance arrays conditioned with SVD, filtered and rectified (Fig. 8).
2. Conditioned arrays transformed into absorbance-like spectra **L**_0_ (Fig. 2AB)

#### Baseline correction

3. Arrays **L**_0_ fitted with exponential relaxations, yielding transient spectra and raw baseline spectra **a, b**_0_ (Fig. 2C).
4. Baseline spectra matched to remove constant effects, **b** (Fig. 2D).
5. Arrays baseline-corrected, **L** = **L**_0_ − **b**_0_, and concatenated, **L**_C_ = [**L**] (Fig. 3A.)

#### Singular value decomposition of concatenations

6. Baseline-corrected arrays **L**_C_ (Fig. 3CD).
7. Transient and matched baseline spectra [**a, b**] (Fig. 4A).
8. Matched baseline spectra [**b**] (Fig. 4A).
9. Transient spectra [**a**] (Fig. 4B).

#### Templates from transient provenience data, all with constraint *r*_*β*M_ ≡ 0

10. Baseline-corrected arrays {**L**}, yielding *α*_R_(*t*), *α*_M_(*t*), used in Step 27.
11. SVD of the concatenated array, **ū**_1_(**L**_C_), Fig. 3;
12. SVD of transient spectra, **ū**_1_([**a**]), Fig. 4.
13. * Set of 21 transient spectra, {**a**}, Fig. 5;

#### From baseline provenience data

14. SVD of baselines, **ū**_1_([**b**]), constraint *r*_*β*M_, Fig. 4;

#### From mixed provenience data

15. SVD of transients and baselines, **ū**_1,2_([**a, b**]), Fig. 4;
16. Same, constraint *r*_*β*R_, Fig. 4;
17. Same, constraint *λ*_*α*R_, *λ*_*α*M_, Fig. 4;

#### From data of both proveniences

18. * Set of transient-baseline pairs, {**a, b**}, Fig. 5;
19. * Same, constraint *r*_*β*R_, Fig. 5;

(* *Datasets providing sample confidence intervals*.)

#### Linear component normalisation

20. Pairs of linear components from Step 13.
21. Quadruplets of linear components from Step 18 (Sec. 3.5).
22. Metarhodopsin to rhodopsin alpha peak ratio, *q*_*α*_ ≈ 1.3 (Fig. 6A).
23. Linear transient and baseline components normalised using *q*_*α*_ (Fig. 6BC).
24. Shutter correction for transient components (Sec. 3.5).
25. Zero-reference for the pigment content (Sec. 3.5).

#### Additional exponential relaxation fits

26. Piecewise fitted SVD component 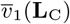 (Fig. 3C).
27. Timecourses *α*_R_(*t*), *α*_M_(*t*) from step 10. (Fig. 7).

## Footnotes

∗ personal communication with Andrej Meglič, University Medical Centre Ljubljana

